# Indirect Sexual Selection Drives Rapid Sperm Protein Evolution

**DOI:** 10.1101/682062

**Authors:** Damien B. Wilburn, Lisa M. Tuttle, Rachel E. Klevit, Willie J. Swanson

## Abstract

Sexual selection can explain rapid evolution of fertilization proteins, yet sperm proteins evolve rapidly even if they are not directly involved in fertilization. Here we demonstrate that FITZAP, an intrinsically disordered sperm protein in the marine mollusk abalone, exploits differences in the intracellular and oceanic ionic environments to package the fertilization protein lysin at extraordinary concentrations inside sperm by forming Fuzzy Interacting Transient Zwitterion (FITZ) complexes. FITZAP binds lysin at the same protein interface as its egg receptor VERL, and as sexual selection rapidly alters the lysin-VERL interface, FITZAP coevolves rapidly to maintain lysin binding. Consequently, FITZAP-lysin interactions exhibit a similar species-specificity as lysin-VERL interactions. Thus, tethered molecular arms races driven by sexual selection can generally explain rapid sperm protein evolution.

**One Sentence Summary:** Structural study of sperm proteins reveals a novel protein packaging/dispersion system embedded in a coevolutionary arms race.

## Main Text

Genes associated with fertilization are often the fastest evolving in any genome (*1*), and in mammals, spermatozoa-specific genes show the greatest divergence between species (*2*). Given the necessity of fertilization for genetic propagation of most animal species, these evolutionary dynamics may seem paradoxical. However, differences in male and female reproductive strategies often result in sexual arms races that drive the rapid evolution of exaggerated sexual phenotypes (*3*). From the elaborate songs of the tungara frog to the ornate plumage of a peacock’s tail, sexual selection on secondary sex characteristics has been hypothesized since Darwin to be a potential driver of speciation (*4–6*). Similarly, sexual selection acting on gamete recognition proteins is postulated to create reproductive barriers and facilitate speciation (*7, 8*), but sexual selection theory does not adequately explain why sperm proteins that do not directly interact with the egg also evolve rapidly. Here we demonstrate that sexual selection can propagate through protein interaction networks and potentially drive global evolution of the sperm proteome.

The marine mollusk abalone is a classic system to study molecular barriers to hybridization (*9*) and is the source of the first discovered pair of interacting reproductive proteins: sperm lysin and the egg <u>v</u>itelline <u>e</u>nvelope <u>r</u>eceptor of <u>l</u>ysin (VERL) (*10*). VERL is a major component of the vitelline envelope (VE), an extracellular barrier that protects the egg and restricts the entry of sperm. Lysin dissolves the VE by binding to repetitive domains within VERL (*10, 11*), and mutations in VERL have resulted in positive sexual selection on lysin that has produced a coevolutionary chase to maintain binding affinity. Consequently, extant lysin proteins dissolve conspecific VEs more efficiently than those of closely related taxa, providing one mechanism of species-specific fertilization and a barrier to hybridization (*10, 12*). The rate of VE dissolution is positively correlated with lysin concentration, and selection has favored sperm that express enormous quantities of lysin (*9*). A single male abalone can contain >1 gram of lysin, reflecting > 0.1% of its total body weight. Within sperm cells, lysin is stored in a specialized secretory granule termed the acrosome. electron microscopy imaging (*13, 14*), we estimate that the acrosomal concentration of lysin is ~0.1 – 1.0 M, in stark contrast to reported saturation concentrations of ~0.001 M under *in vitro* conditions (*15*). We therefore investigated what special conditions permit such extraordinarily high lysin concentrations within the sperm acrosome.

Lysin is not the only highly abundant, rapidly evolving protein in the abalone sperm acrosome. Another is sp18, a fusogenic paralog of lysin that likely mediates plasma membrane fusion between egg and sperm (*16*). The receptor of sp18 is unknown, but given its interaction with the abalone egg, its accelerated evolution is likely due to sexual selection (*17*). Sp18 is nearly as abundant as lysin, so it must also be packaged at high concentrations, yet its fusogenic properties make it even less soluble than lysin (*18*). Recently, a new family of small acrosomal proteins termed sperm protein 6 kDa (sp6) was discovered by shotgun transcriptomics and proteomics (*19*). While also rapidly evolving and hypothesized to evolve via sexual selection, initial efforts to identify an egg binding partner for sp6 were unsuccessful (*20*). However, while lysin and sp18 are highly positively charged proteins (+12 to +24), isoforms of sp6 are highly anionic (−6 to −16) and include an N-terminal poly-aspartate region of variable length (1 – 11 residues). Given this charge complementarity, we hypothesized that sp6 may facilitate packaging of lysin and sp18 inside the sperm acrosome. We demonstrate that the rapid evolution of sp6 is due to intra-sperm protein coevolution with lysin and sp18 to allow for their dense storage in the acrosome via novel Fuzzy Interacting Transient Zwitterion (FITZ) complexes. Heterodimers of lysin-sp6 or sp18-sp6 form through hydrophobic interactions, and these heterodimers polymerize into large particles (diameter > 100 nm) through ionic interactions of the complementary positive and negative charges. Upon secretion of the acrosomal contents into highly ionic seawater, FITZ complexes are disrupted, with lysin and sp18 dispersed to facilitate fertilization. In light of its newly identified function, we have named sp6 the FITZ Anionic Partner (FITZAP).

Different species of abalone express different numbers of FITZAP isoforms named for the length of the N-terminal poly-aspartate region (*20*). In red abalone (*Haliotis rufescens*), two isoforms (FITZAP-4D and FITZAP-8D) result from alternative splicing of two different versions of exon 1 (that contains a signal peptide and the N-terminus with the poly-aspartate region) with a common exon 2 (that contains the majority of the C-terminus) (Fig S1). Through a combination of strong anion exchange (SAX) chromatography and reverse-phase high performance liquid chromatography (RP-HPLC), each isoform was purified to homogeneity. Mass spectral analysis revealed that both isoforms were smaller than their cDNA opening reading frame predicted (~3-4 kDa vs ~6 kDa). The observed masses are consistent with proteolytic processing of FITZAP in the Golgi complex by the proteases furin and carboxypeptidase B (Fig S2). Despite their high net positive charges, both lysin and sp18 co-eluted with FITZAP at high salt concentrations by SAX chromatography (Fig S3). Particularly striking is that sp18 (+22) eluted at higher salt concentrations than lysin (+12), which closely mirror the elution profiles of FITZAP-4D (−10) FITZAP-8D (−8), respectively, suggesting isoform-specific interactions. While purified lysin showed no affinity for anion exchange resin, its elution was retarded when mixed with either FITZAP-4D or FITZAP-8D *in vitro*, consistent with the co-elution of lysin/sp18 and FITZAP from sperm lysate resulting from *in vivo* interactions.

Nuclear magnetic resonance (NMR) spectroscopy was used to investigate how FITZAP interacts with the cationic fertilization proteins. We focused on lysin and FITZAP-8D because (A) they showed the strongest coelution by SAX chromatography (Fig S3B), (B) lysin is more soluble than sp18 (*16, 17*), (C) interactions with its egg receptor have been characterized (*11, 15*), and (D) a solution structure has been determined (*15*). Chemical shift analysis of FITZAP-8D revealed that it is an intrinsically disordered protein (IDP); even when bound to lysin, FITZAP-8D remained highly dynamic and adopted no regular secondary structure (Fig S4). Thus, lysin and FITZAP form a fuzzy complex: protein complexes that exist in an ensemble of different interchanging configurations. Formation of the fuzzy complex is primarily due to packing between hydrophobic amino acids in residues 21-29 of FITZAP (Fig S5) and an exposed hydrophobic face on lysin near the nexus of the N- and C-termini. Covering this hydrophobic region with FITZAP imparts a high net negative charge to this face of lysin, likely explaining the affinity of lysin to anion exchange resins when FITZAP is present. Significantly, the FITZAP-binding region of lysin is also the same surface that recognizes its egg receptor VERL (Fig 1). We postulate that this exposed hydrophobic surface is responsible for purified lysin having a much lower *in vitro* solubility limit compared to the acrosome where FITZAP is also highly abundant.

**Fig. 1.**
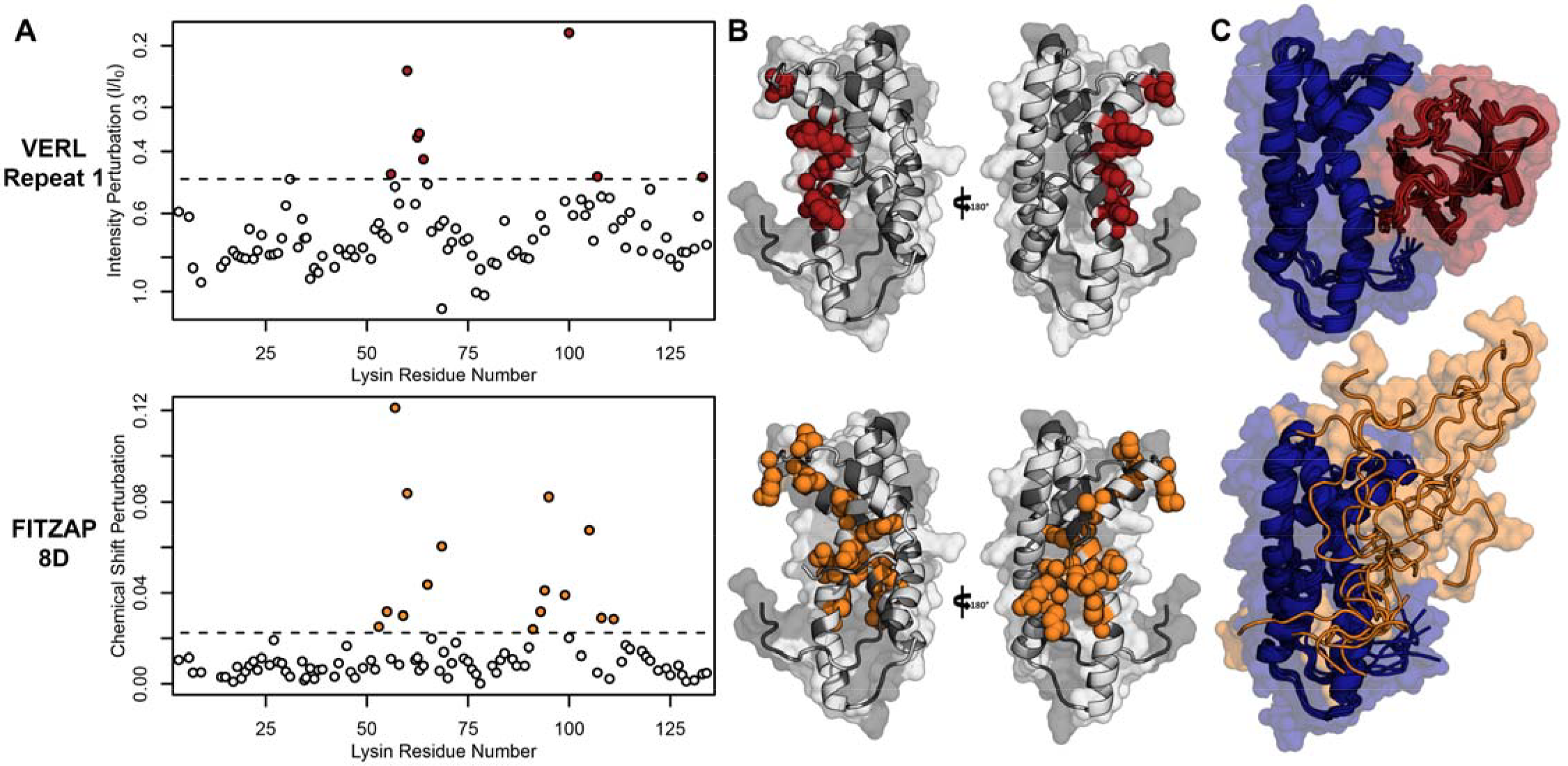
Lysin recognizes VERL and FITZAP through a common binding interface. (A) NMR perturbation of red lysin upon conspecific binding of either VERL repeat 1 (data from Wilburn et al. (*15*)) or FITZAP-8D. Two regions near residue 60 and residue 100 of lysin are perturbed in both cases (dashed line: median + 1.5 * interquartile range). (B) Mapping of perturbation onto a lysin solution structure (PDB 5utg) shows spatial clustering of these two regions to a single binding surface of lysin. (C) Docking of VERL repeat 1 (based on PDB 5mr3 by Raj et al. (*11*)) and FITZAP-8D (based on restraints from paramagnetic relaxation enhancement) supports that FITZAP is an intrinsically disordered protein that shields the VERL binding interface of lysin.

Given a single interaction surface for two different identified binding partners, lysin must separately recognize FITZAP inside the acrosome and then VERL once it is secreted upon contact with the VE. As abalone is a marine mollusk, we postulated that local inorganic salt concentrations may play an important role. The total salt concentration of seawater is ~500 mM, yet the intracellular environment of marine animals is less concentrated. In fish and marine mammals osmolality is principally maintained by active transport of water and ions; instead, abalone are osmoconformers where osmolality (but not isotonicity) is established by high intracellular concentrations of free amino acids, betaines, and other highly soluble metabolites (*21*). While the exact intracellular concentration of inorganic ions in abalone sperm is unknown, it is likely much lower than seawater based on data from sea urchin gametes (*22, 23*) and other abalone tissues (*24*). The inorganic salt concentration may be even lower in the acrosome where – given their extraordinary concentrations and net charges – the acrosomal proteins may themselves serve as osmolytes. To compare how lysin-FITZAP interactions may be influenced by differing environmental contexts, we performed biophysical experiments under low (150 mM NaCl) and high (500 mM NaCl) salt concentrations approximate the intracellular or seawater environments, respectively. Under both conditions, the hydrophobic patch of FITZAP is found to interact with lysin, yet NMR intensity perturbation of the poly-aspartate region was only observed under low salt concentrations where they may be forming intermolecular salt bridges (Fig S5). Consistent with additional types of molecular interactions, differences in NMR perturbation establish that lysin and FITZAP undergo slower subunit exchange under low salt concentrations (Fig S6). Equimolar mixtures of lysin and FITZAP-8D form extremely large oligomers with an average diameter of ~400 nm (compared to a mean lysin diameter of ~6 nm) under low salt conditions; these large particles are not present at high salt concentrations (Fig 2). Altogether, our data establish that lysin and FITZAP associate hydrophobically to form fast-exchanging, fuzzy heterodimers that, although highly charged, are essentially zwitterionic. These heterodimers that we call <u>F</u>uzzy <u>I</u>nteracting <u>T</u>ransient <u>Z</u>witterions (FITZs) can form intermolecular salt bridges that allow tight packaging under intracellular-like conditions. Upon secretion into seawater when sperm contact an egg, the FITZ complexes are disrupted and subunit exchange rate increases, allowing lysin to be rapidly liberated from FITZAP, permitting interactions with VERL via a common binding interface. The formation and dissolution of FITZ complexes based on the environmental context provides a mechanism for exceptionally high packaging concentrations of lysin inside the sperm acrosome as well as its rapid dispersal in seawater when fertilization may be imminent.

**Fig. 2.**
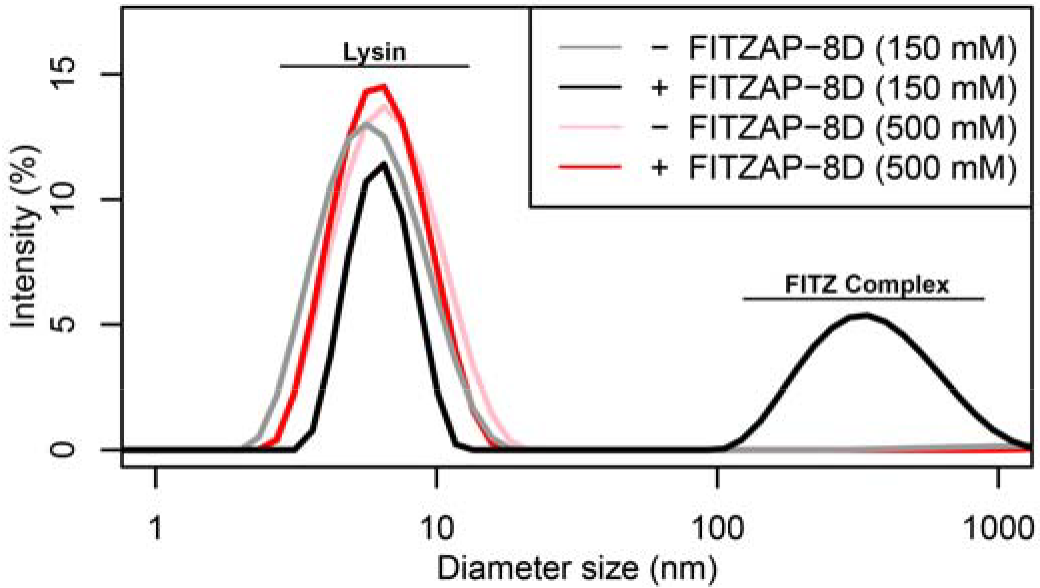
FITZ complex formation is dependent on both FITZAP and the ionic environment. Dynamic light scattering measurements of lysin with and without equimolar FITZAP-8D at intracellular (150 mM) and seawater (500 mM) salt conditions. Lysin and FITZAP associate into FITZ complexes with mean diameter of ~ 400 nm only under intracellular salt conditions.

Lysin and VERL are rapidly coevolving sperm and egg proteins that exhibit species-specific interactions (*25*). Given the common binding interface, we postulated that there may be similar coevolution between lysin and FITZAP. While coevolution between lysin and FITZAP was not detected using sequence-based approaches (Fig S7), the small size and intrinsic disorder of FITZAP can reduce statistical power of such analyses. A functional consequence of lysin and FITZAP coevolution would be species-specific interaction, as observed between lysin and VERL. Therefore, lysin and FITZAP from the same species should have higher binding affinity compared to heterospecific pairs. Using fluorescence polarization, binding affinities were measured between lysin and all FITZAP isoforms for three abalone species (red, disk, and green). Like red abalone, green abalone has two FITZAP isoforms but with shorter poly-aspartate regions (1D and 4D), while disk abalone has a single FITZAP isoform with an even longer poly-aspartate array (11D). Lysin had greater affinity for the high-D FITZAP isoforms compared to low-D forms. For all three species, lysin bound the conspecific high-D isoforms of FITZAP with equilibrium dissociation constants of ~1-2 μM under intracellular salt conditions. Except for disk lysin and red FITZAP-8D, all cases of heterospecific binding were significantly weaker, consistent with lysin-FITZAP coevolution (Fig 3A). Under seawater conditions, there was a ≥ 20-fold decrease in conspecific binding affinity (Fig 3B), consistent with the liberation of lysin from FITZ complexes after its release from the acrosome.

**Fig. 3.**
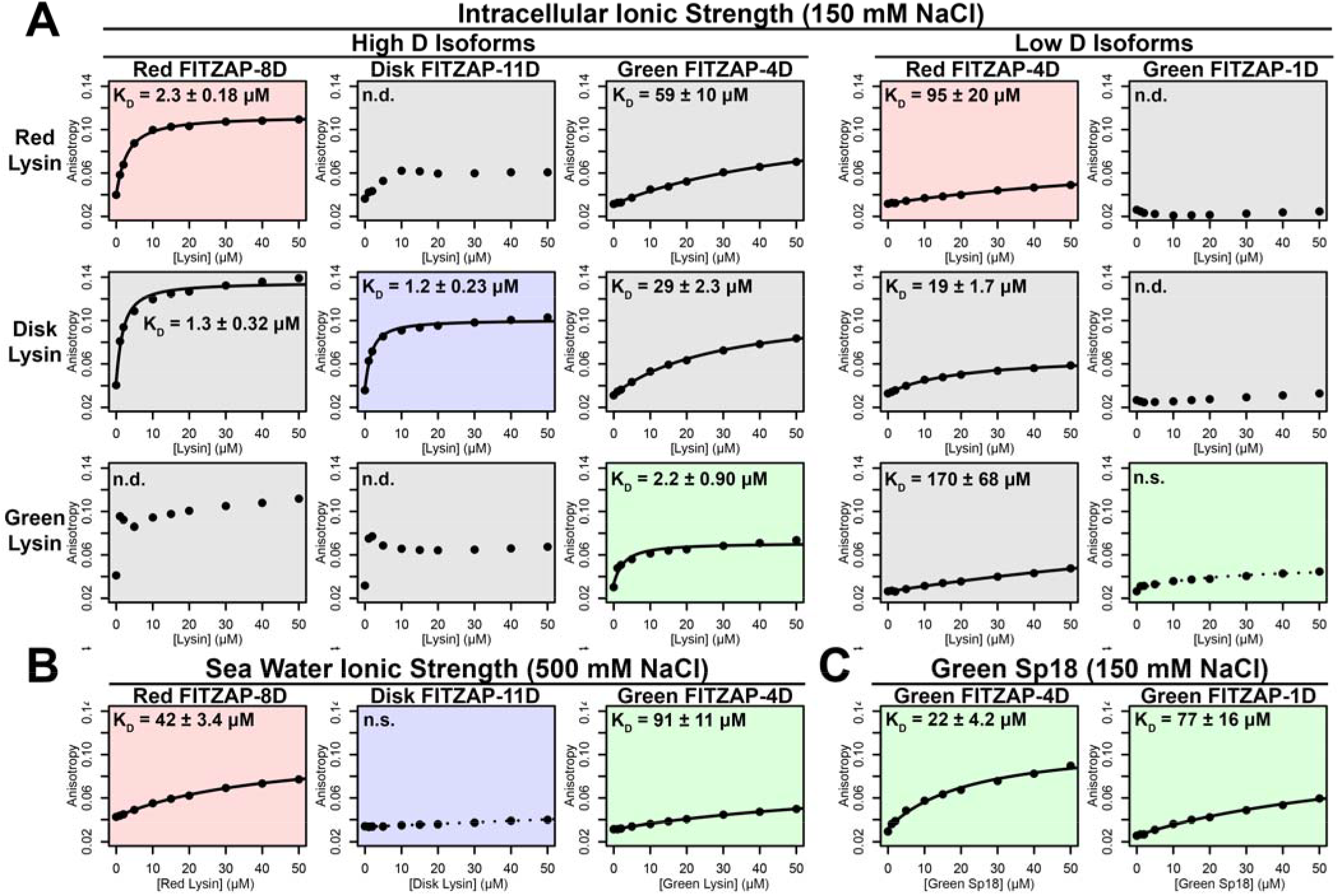
FITZAP interactions with cationic acrosomal proteins within and between species. (A) Lysin-FITZAP interactions were measured by fluorescence polarization under approximately intracellular salt conditions for all combinations within and between species of red, disk, and green abalone (conspecific interactions are shaded as red, blue, and green, respectively). Except in the case of disk lysin with red FITZAP-8D, low micromolar binding affinities were observed for conspecific interactions of lysin with high-D FITZAP isoforms. (B) Binding affinities between conspecific lysin and FITZAP high-D isoforms are substantially reduced under extracellular seawater conditions. (C) Green sp18 shows tighter binding to conspecific FITZAP-4D compared to FITZAP-1D; however, the relative affinity of sp18 for 4D over 1D (77/22= 3.5 X) is relatively greater than that of lysin (> 100 X), suggesting that sp18 has relatively higher preference for low D isoforms compared to lysin. (n.s. = not significant at p < 0.05; n.d. = not determined if anisotropy was not monotonically positive consistent with single-state binding).

We propose a system of tethered coevolution between VERL, lysin, and FITZAP where VERL imposes direct sexual selection on lysin and indirect sexual selection on FITZAP (summarized in Fig 4). As with any coevolving system, there is likely reciprocity with all binding partners imposing some form of selection on one another; however, we choose to focus on the unidirectional case of VERL influencing lysin influencing FITZAP for several reasons. First, sexual selection theory has classically emphasized “female choice,” because the higher energetic cost of oocytes in most species should favor greater mate selectivity (*7*). These assumptions are most apt when discussing genes involved in fertilization. Second, lysin evolves ~5X faster than its coevolving regions of VERL (*26*), suggesting that it is experiencing greater directional selection than VERL. Lysin, but not VERL, is also monomorphic within some abalone populations and experienced recent selective sweeps (*27*). Third, in sea urchins (another organism with broadcast spawning), longitudinal measurements of allele frequencies for gamete recognition proteins support that female proteins will shift to lower affinity interactions in response to increased polyspermy risk (positive natural selection), which is then followed by male proteins adapting to restore high affinity binding (positive sexual selection) (*28*). Together, this supports that evolutionary dynamics of lysin are more responsive to VERL than vice versa. Fourth, as FITZAP is an IDP whose function is to facilitate the storage and dispersal of cationic acrosomal proteins, we anticipate minimal selection on its primary sequence beyond directional selection imposed by its binding partners.

**Fig. 4.**
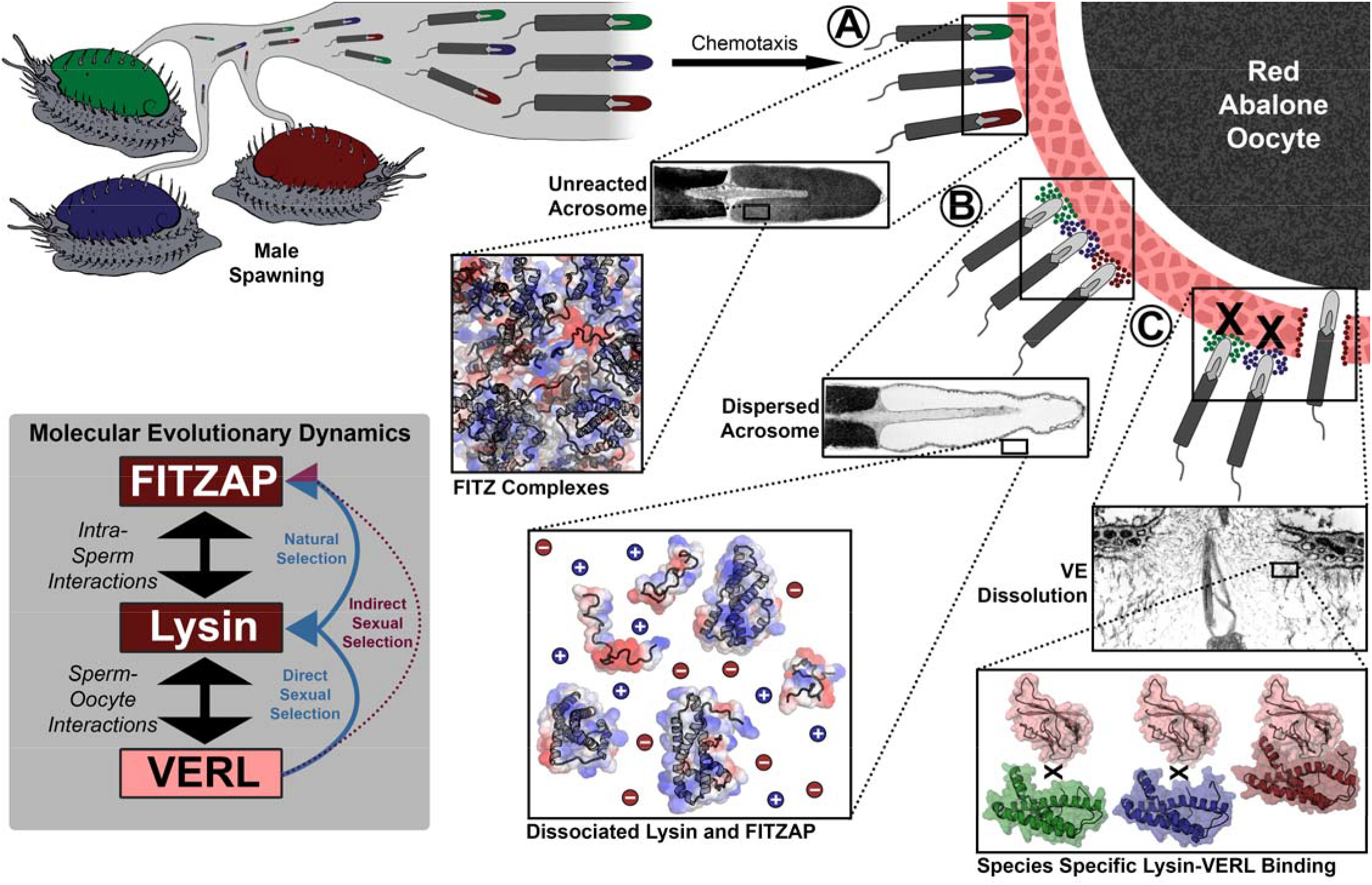
Rapid evolution of FITZAP is due to indirect sexual selection from VERL. Male abalone have overlapping habitat ranges and spawning periods such that there is opportunity for hybridization to occur. Sperm are attracted to eggs via chemotaxis (with L-tryptophan as the chemoattractant) (*29*), sperm bind to the vitelline envelope **(A)**. At this stage, the acrosome is still intact (*9*) with lysin and FITZAP tightly packaged through electrostatic FITZ complex interactions. Binding to the VE causes the sperm to acrosome react **(B)**, releasing its contents into highly ionic seawater which disperses lysin and FITZAP. Liberated lysin can now dissolve the VE by binding VERL domains **(C)**; these interactions are species-specific and provide one barrier to hybridization (*10*). Because the rapid evolution of lysin is driven by direct sexual selection to maintain sperm-egg interactions, and FITZAP is co-evolving to maintain affinity to the same interface that is evolving via direct sexual selection, VERL imposes indirect sexual selection onto FITZAP. Microscopy images adapted from Lewis et al. (*9*) and Sakai et al. (*30*).

Because IDPs lack a single favored conformation, they generally experience less purifying selection to maintain a tertiary fold and have lower sequence conservation (*31*). Beyond relaxed purifying selection and greater rates of genetic drift, IDPs also more commonly experience positive selection compared to structural domains (*32*), presumably in response to coevolution with their binding partners. As such, we expect FITZAP to respond more to selection from lysin than vice versa. It has been suggested in the literature that intrinsic disorder may be a mechanism of adaptation to shifts in environmental conditions; for example, host-changing parasites have higher genome-wide levels of predicted protein disorder compared to obligate intracellular parasites and endosymbiotes (*33*). The change in ionic strength experienced by secreted proteins of most marine animals is likely an extreme example of such shifts between chemical environments. Thus, the intrinsic disorder of FITZAP may be crucial to its apparent structural versatility and high evolvability in response to a rapidly coevolving partner.

As sp18 likely facilitates the fusion of egg and sperm plasma membranes (*16*), it is even more hydrophobic than lysin and may therefore be even more reliant on a partner for storage and dispersal. Lysin and sp18 are paralogs with similar tertiary structures that have subfunctionalized following gene duplication (*16, 18*), so the multiple FITZAP isoforms may also have subfunctionalized to serve these different paralogs. As lysin showed greater affinity for high-D FITZAP isoforms (Fig 3A), we hypothesized that sp18 may be coevolving with low-D isoforms. Across Pacific abalone, we indeed observe correlated rates of molecular evolution between sp18 and low-D FITZAP isoforms, consistent with coevolution (Fig S7). While the highly fusogenic sp18 is mostly insoluble when purified (*16, 17*), we were able to measure conspecific affinities between sp18 and FITZAP isoforms from green abalone. While green sp18 also bound green FITZAP-4D more tightly than FITZAP-1D (Fig 3C), its relative affinity for 4D over 1D (~3.5X) is substantially less compared to the relative affinity of 4D to 1D for lysin (>100X). Additionally, for our anion exchange experiments in red abalone, we observed tighter co-elution between lysin/FITZAP-8D and sp18/FITZAP-4D (Fig S3). Therefore, both evolutionary and biochemical analyses support that FITZAP isoforms have subfunctionalized to respond to the divergent evolutionary trajectories of lysin and sp18.

Across animals, plants, and microbes, genes associated with fertilization usually evolve faster than the rest of the genome (*1, 8, 25*). Like macroscopic secondary sex characters, direct sexual selection can drive elaboration of molecular phenotypes such as the extraordinary abundance of lysin and fusogenicity of sp18. Combined with sequence differences that yield species-specific interactions with coevolving receptors, these proteins are one of many reproductive barriers that likely contribute to speciation. Given the necessity of fertilization for sexually reproducing taxa, few selective pressures are likely stronger than sexual selection, and even its indirect effects through coevolutionary networks are likely substantial. In this report, we demonstrated how indirect sexual selection drives rapid evolution of a sperm protein not associated with fertilization. Hence, we postulate that such indirect selection may further propagate throughout the sperm proteome and be a general mechanism to explain accelerated gametic protein evolution.

## Acknowledgements

The authors would like to thank Jan Aagaard, Emily Killingbeck, Jolie Carlisle, Alberto Rivera, Richard Feldhoff, Auberon Lopez, and Harmit Malik for feedback on the research and/or manuscript.

## Author Contributions

The study was conceived and designed by DBW and WJS. All experiments and data analyses were performed by DBW and LMT. The manuscript drafted by DBW and edited by LMT, REK, and WJS.

## Data and materials availability

Sequence data has been deposited into Genbank (Accession # MN102340-MN102343) and NMR assignments have been deposited to the BMRB (27962). All other data is available in the main text or supplementary materials.

