## Supplemental Material for "Indirect Sexual Selection Drives Rapid Sperm Protein Evolution"

**This PDF file includes:**

Materials and Methods
Figs. S1 to S7

### Materials and Methods

*Purification and mass spectral characterization of natural FITZAP*

Natural FITZAP was purified and characterized based on methods modified from Palmer et al. (*19*). Briefly, sperm were collected by dissection of testes from red abalone (*Haliotis rufescens*) and lysed by trituration in 1% Triton X-100 (w/v)/250 mM NaCl/2 mM EDTA/10 mM MES, pH 6, and centrifuged at 3200 *x g*, 8°C for 30 minutes. The supernatant was applied to a 10 mL CM52 cellulose column (Whatman, Maidstone, UK) to remove lysin and other positively charged proteins. The FITZAP-enriched flow through was diluted with 2 volumes of 50 mM Tris, pH 8 and applied to a 10 mL Q sepharose column (Sigma-Aldrich, St. Louis, MO) and protein fractions collected by gravity flow with a stepwise NaCl gradient buffered with 20 mM Tris, pH 8. Fractions were analyzed by 15% Tris-Tricine SDS-PAGE (*34*), with FITZAP localized to fractions with ≥ 300 mM NaCl. These fractions were pooled and concentrated using 3 kDa centrifugal ultrafilters (EMD-Millipore, Billerica, MA), and individual components purified by reverse phase high performance liquid chromatography (RP-HPLC) using a Vydac C18 column (0.46 x 15 cm; Hichrom, Berkshire, UK) that was eluted from 0 – 70% acetonitrile in 0.1% trifluoracetic acid at 1% acetonitrile per minute. Individually purified proteins were analyzed by LC/MS-MS using data-dependent acquisition on an LTQ Velos tandem mass spectrometer (Thermo Scientific, Waltham, MA) for determination of intact protein and fragment ion masses.

*Sequence analysis of FITZAP and estimation of molecular evolutionary rates*

Draft genome assemblies are available for disk abalone (*Haliotis discus*) (*35*) and red abalone (*H. rufescens*) (*36*). The contigs or scaffolds containing FITZAP exons were identified by performing BLAST searches (*37*) using the FITZAP open reading frames as queries. These genomic regions were extracted with an additional 5 kb of flanking sequence on both the 5' and 3' ends, and aligned using fsa v1.15.9 (*38*). For molecular evolutionary analysis, available cDNA sequences for lysin, VERL, sp18, and FITZAP were downloaded from Genbank (Accession # M34388, M59969-M59972, M98875, AF453553, AF490761-AF490763, AF490765, AF490766, L36552, L36554, L36589, AGJ90053, AGJ90054, AGJ90056-AGJ90061). We additionally sequenced sp18 cDNA by RT-PCR from black abalone (*Haliotis cracherodii*), flat abalone (*Haliotis wallallensis*), and disk abalone (*H. discus*), and FITZAP from white abalone. Abalone testis cDNA was provided by Jan Aagaard, and RT-PCR was performed using an oligo-dT reverse primer using a sp18 specific (5'-GGAAACAGTATGAGGTYTTTGSTGCTT-3') or FITZAP-specific (5'-ATGAGGGTTRTTCTAATT-3') forward primer. PCR products were cloned into a pCR4-TOPO vector (Invitrogen), transformed into 5-alpha competent *E. coli* (New England Biolabs, Ipswich, MA), and plasmid DNA from at least 4 clones of each transformation were supplied to Eurofins Scientific (Louisville, KY) for Sanger sequencing. No sequence variation was observed between clones, and these sequences have been deposited into Genbank (Accession # MN102340-MN102343). A maximum likelihood gene tree was constructed with RAxML v8.2.12 (*39*) using the PROTGAMMALG substitution model and a concatenation of protein sequences from lysin, VERL, and sp18 from 6 abalone species (red, flat, disk, black, pinto (*Haliotis sorenseni*), and green (*Haliotis fulgens*)) aligned by Clustal Omega (*40*), and used as a representative of the likely species tree. Based on methods by Clark et al. (*27*), support for co-evolution between protein coding genes was evaluated by weighted linear regression using branch dN/dS values estimated with PAML v4 (*41*) for each of the four genes (lysin, VERL, sp18, and FITZAP), with FITZAP further divided into low-D and high-D isoforms.

*Cloning and expression of recombinant FITZAP*

Recombinant FITZAP was expressed in *E. coli* and purified to near homogeneity using multiple chromatography steps. Given the low molecular weight of different FITZAP isoforms, expression in *E. coli* required fusion to a larger carrier protein from which FITZAP could be removed by enzymatic proteolysis and purified. FITZAP isoforms from red, disk, and green abalone were genetically fused by PCR with a maltose binding protein (MBP) cassette containing an N-terminal 6xHis tag and a C-terminal linker sequence followed by a tobacco etch virus (TEV) protease cleavage site. Recombinant FITZAP proteins included an Amino Terminal Cu- and Ni-binding tag (ATCUN) to facilitate protein purification, improve TEV proteolysis (*42*), and permit collection of NMR paramagnetic relaxation enhancement (PRE) constraints (*43*). The combined MBP-FITZAP construct was cloned into the pET11d expression vector (Novagen, San Diego, CA), transformed into Rosetta2 chemically competent *E. coli* (EMD-Millipore, Billerica, MA) which express additional tRNA genes for Lys and Arg that are abundant in abalone genes, and clones validated by Sanger sequencing (Eurofin Genomics, Louisville, KY). For expression of unlabeled FITZAP, *E. coli* clones were cultured in LB media supplemented with 100 µg/mL ampicillin and 34 µg/mL chloramphenicol at 37°C, 250 rpm; when cultures reached an optical density at 600 nm (OD600) of ~0.6, recombinant protein expression was induced by addition of IPTG to a final concentration of ~100 µM, cells harvested by centrifugation after 3.5 hours, and stored at -20°C. For expression of isotopically labeled FITZAP, methods were adapted from Wilburn et al. (*15*). Briefly, *E. coli* clones expressing MBP-FITZAP were cultured in LB media supplemented with 100 µg/mL ampicillin and 34 µg/mL chloramphenicol at 37°C, 250 rpm until the OD600 reached ~0.4; cells were then collected by centrifugation, concentrated 4-fold into M9 media with 20 µM $FeSO_4$ and 100 µg/mL ampicillin without carbon or nitrogen sources, and maintained at 37°C, 250 rpm for 35 min to deplete the cells of free unlabeled amino acids; cultures were then supplemented with 3 g/L ammonium sulfate ($^{14}N$ or 98% $^{15}N$) and 4 g/L glucose ($^{12}C$ or 99% $^{13}C$) for 35 min to regenerate amino acid stores with appropriate isotopes; then expression was induced by addition of IPTG to a final concentration of ~100 µM, cells harvested by centrifugation after 3.5 hours, and stored at -20°C. Cell pellets were then lysed by sonication in 1% octylthioglucoside/50 mM NaCl/50 mM Tris, pH 8, then supplemented with 0.2 mg/mL lysozyme for 30 min, centrifuged @ 3.2k *x g* for 2 hours, the supernatant clarified by passage through a 0.2 µm PES filter, and the filtrate applied to a 10 mL Ni-NTA column (Pierce, Rockford, IL) equilibrated in 500 mM NaCl/20 mM Tris/1 mM imidazole, pH 8. The column was subsequently washed with increasing concentrations of imidazole in 500 mM NaCl/20 mM Tris, pH 8: 6 column volumes (CVs) at 1 mM imidazole, 2 CVs at 20 mM, 1 CV at 40 mM, 1 CV at 60 mM. and MBP-FITZAP eluted using 3 CVs at 200 mM imidazole. The elution fraction was buffer exchanged using a YM30 centrifugal ultrafilter (Millipore, Billerica, MA) into 100 mM NaCl/20 mM Tris, pH 8, supplemented with TEV Protease (Sigma-Aldrich) at an enzyme:substrate ratio of ~1:500 by mass, and incubated overnight at room temperature with gentle mixing. Following proteolysis, precipitate was removed by centrifugation at 3.2k *x g* for 30 min, and the supernatant applied to a 10 mL Ni-NTA column equilibrated in 500 mM NaCl/20 mM Tris/1 mM imidazole, pH 8. The column was washed with 3 CVs of 500 mM NaCl/20 mM Tris/1 mM imidazole, pH 8, a FITZAP-enriched fraction eluted using 3 CVs of 500 mM NaCl/20 mM Tris/20 mM imidazole, pH 8, and MBP/MBP-FITZAP enriched fraction eluted using 500 mM NaCl/200 mM Tris/1 mM imidazole, pH 8. The 20 mM imidazole FITZAP-enriched fraction was further purified by size-exclusion chromatography (G-75 superfine; Pharmacia, Piscataway, NJ) followed by strong anion exchange chromatography (Mono-Q; Pharmacia) on an Agilent 1100 HPLC with UV detection at 220 nm.

*Recombinant lysin expression*

Recombinant lysin from red and disk abalone was expressed and purified by methods from Wilburn et al. (*15*). Briefly, lysin coding sequences were cloned into the pET11d expression vector (Novagen), transformed into Rosetta2 chemically competent *E. coli* (EMD-Millipore, Billerica, MA) which express additional tRNA genes for Lys and Arg that are essential for lysin expression, and clones validated by Sanger sequencing (Eurofin Genomics, Louisville, KY). To provide flexibility in isotopic labeling for NMR experiments, lysin expression was performed in cultures where (1) biomass with high ribosome densities was produced by initially culturing in complex media, (2) the cells were concentrated ~4X in minimal media without nitrogen or carbon to deplete amino acid stores, (3) ammonium sulfate ($^{14}N$ or $^{15}N$) and glucose (uniformly $^{12}C$ or $^{13}C$) were added to regenerate amino acids with the appropriate isotopes, then (4) expression was induced by addition of IPTG. Growth under minimal media conditions provides complete removal of the N-terminal methionine from endogenous *E. coli* methionine aminopeptidase activity, leaving a single exogenous Gly on the N-terminus that has no detectable impact on lysin structure or function. Properly folded recombinant lysin was expressed into inclusion bodies that were isolated by centrifugation of cell lysate, washed to remove contaminant proteins, denatured in 5M guanidinium hydrochloride, refolded by rapid dilution, and purified using cation exchange chromatography.

*Ion exchange purification of lysin and sp18*

Methods for lysin purification were adapted from Lewis et al. (*9*). For both natural and recombinant lysin, step chromatography was performed using CM52 cellulose (Whatman) equilibrated in 250 mM NaCl/10 mM MES/2 mM EDTA, pH 6. For recombinant lysin, methods are described above for refolding from inclusion bodies. For natural lysin, *H. rufescens* sperm were isolated by dissection of male testes and lysed by trituration in 0.1% Triton X-100/250 mM NaCl/10mM MES/2mM EDTA, pH 6; insoluble material (including chromatin) was removed by centrifugation at 3,200 *x g* for 30 minutes. Crude fractions of natural or recombinant lysin were applied to a CM52 cellulose column, rinsed with > 6 CVs of 250 mM NaCl/10 mM MES/2 mM EDTA, pH 6, and eluted with 3 CVs of 1 M NaCl/10 mM MES/2 mM EDTA, pH 6. Purified natural lysin and sp18 from *H. fulgens* was generously supplied by Vic Vacquier. Purified lysin and sp18 was concentrated and buffer exchanged to 150 mM NaCl/10 mM Tris, pH 7.4 using YM10 centrifugal ultrafilters (Millipore).

*Comparison of conspecific/heterospecific FITZAP-Lysin/Sp18 interactions by fluorescence polarization*

Purified recombinant FITZAP proteins from red, disk, and green abalone were buffer exchanged into 0.5 mL phosphate buffered saline (PBS) using a 3 kDa centrifugal ultrafilter (Millipore), and fluorescently labeled by addition of 18 µL Alexa Fluor 488 SDP (Invitrogen, Carlsbad, CA) at 10 mg/mL DMSO at 4°C overnight with mixing. Fluorescently labeled FITZAP was separated from free fluorophore by size exclusion using a Nap5 column (GE Life Sciences, Piscataway, NJ). Protein concentrations for labeled FITZAP and unlabeled lysin were determined by BCA Protein Assay (Pierce). Each fluorescently labeled FITZAP isoform was standardized to 1 µM, lysin added to concentrations of 0, 1, 2, 5, 10, 15, 20, 30, 40, and 50 µM, and fluorescence anisotropy measured using a Fluorolog spectrofluorometer (Horiba Scientific, North Edison, NJ). All species/isoform combinations between FITZAP and lysin were performed in 150 mM NaCl/10 mM Tris, pH 7.4, and when possible, anisotropy measurements were collected in technical duplicate (although sample degradation over the course of the experiment prevented this for all combinations). For conspecific pairings with low µM binding affinities, anisotropy measurements were repeated in 500 mM NaCl/10 mM Tris, pH 7.4. Additionally, anisotropy experiments were repeated for green FITZAP isoforms with green sp18 using the same series of concentrations as lysin in 150 mM NaCl/20 mM Tris, pH 7.4. Dissociation constants ($K_D$) were estimated for all combinations by nonlinear regression using the equation $Anisotropy \sim \Delta A_{max} * \frac{([Lysin]+[FITZAP]+K_d-\sqrt{([Lysin]+[FITZAP]+K_d)^2-4*[Lysin]*[FITZAP]})}{2} + A_{intercept}$ with the R function nlsLM in the package minpack.lm.

*Anion exchange analysis of positively charged acrosomal proteins*

Preliminary experiments separating *H. rufescens* sperm lysate by anion exchange chromatography yielded the surprising result of both lysin and sp18 (highly positively charged proteins) adhering to the column and eluting at relatively high ionic strengths, and it was hypothesized that this unexpected observation may result from FITZAP supplying negative charges to these proteins as part of FITZ complexes at low ionic strength. Sperm from *H. rufescens* was isolated by dissection of testes, lysed by trituration in 2 mL of 0.1% Triton X-100/20mM Tris, pH 8 supplemented with TURBO DNase (Ambion, INFO), and centrifuged at 2k *x g* for 10 minutes. Clarified lysate was applied to a 5 mL Q Sepharose (Sigma-Aldrich) column, and 2 CV fractions collected at 0, 25, 50, 100, 200, 300, 400, and 600 mM NaCl in 20 mM Tris, pH 8. The same step gradient was performed after applying 2 mL aliquots of (1) red lysin at 0.5 mg/mL, (2) red lysin at 0.5 mg/mL with equimolar red FITZAP-8D, and (3) red lysin at 0.5 mg/mL with equimolar red FITZAP-4D. Anionic exchange fractions were separated by 15% Tris-Tricine SDS-PAGE (*34*) and stained with Coomassie Brilliant Blue R-250.

*NMR analysis of FITZAP-Lysin interactions*

Purified, isotopically labeled *H. rufescens* FITZAP-8D ($^{15}N/^{13}C$) was concentrated to ~0.2-1.0 mM in 50 mM NaCl/10 mM Tris, pH 7.4/7% $D_2O$ using a 3 kDa centrifugal ultrafilter (Millipore). For FITZAP, all NMR experiments were performed on a Bruker Avance 800-Mhz spectrometer fitted with a TCI CryoProbe (Bruker). NMR assignments of red FITZAP-8D were obtained using a combination of 2D/3D experiments: $^{15}N$- and $^{13}C$-filtered HSQC, HNCACB, CBCAcoNH, HNCO, HNHA, $^{15}N$-HSQC-TOCSY, and $^{15}N$-HSQC-NOESY. Spectra were processed using NMRpipe (*44*) and analyzed using NMRFAM-SPARKY (*45*). Assignments were 70% complete for backbone atoms (91% excluding the poly-aspartate region). Chemical shift indices were calculated using TALOS-N (*46*). To characterize lysin binding residues, $^{15}N$- and $^{13}C$-HSQC spectra of $^{15}N/^{13}C$-FITZAP-8D (200 μM in 500 mM NaCl/10 mM Tris, pH 7.4/7% $D_2O$) were acquired at 6 concentrations of recombinant monomeric lysin (*15*) from 0 to 500 μM. Chemical shift perturbations (CSPs) between $^{15}N$- and $^{13}C$-HSQC spectra were calculated as $\sqrt{(\Delta\ ^{1}H)^2 + (0.1 * \Delta^{15}N)^2}$ or $\sqrt{(\Delta\ ^{1}H)^2 + (0.1 * \Delta^{13}C)^2}$. The interaction of lysin and salt with FITZAP was assessed by acquiring $^{15}N$- and $^{13}C$-HSQC spectra of $^{15}N/^{13}C$-FITZAP-8D (200 μM in 10 mM Tris, pH 7.4/7% $D_2O$) with or without monomeric lysin (140 μM) at different salt concentrations (150 – 500 mM in 50 mM steps). Reciprocal titration experiments were performed with $^{15}N$- and $^{13}C$-HSQC spectra acquired for $^{15}N/^{13}C$-monomeric lysin (150-200 μM in 10 mM Tris, pH 7.4/7% $D_2O$) at different salt concentrations (150 or 500 mM) with varying concentrations of FITZAP-8D (0 to 600 μM). To obtain intermolecular PRE constraints from the FITZAP-8D N-terminal ATCUN motif, R2 relaxation rates were measured for $^{15}N$- monomeric lysin (150 μM in 500 mM NaCl/10mM Tris, pH 7.4/7% $D_2O$) with equimolar FITZAP-8D with or without 135 μM $CuSO_4$ using delays of 8.48, 16.96, 25.44, 33.92, 42.40, 50.88, and 59.36 ms. Data has been deposited in the BMRB (27962).

*Structural analysis*

Using Xplor-NIH 2.48 (*47, 48*), a structural ensemble of lysin and FITZAP-8D heterodimers was modeled by simulated annealing from 4000 to 25 K with torsion dynamics followed by Cartesian minimization using constraints from PRE measurements and degenerate CSP pairings (adapted from Clore and Schwieters (*49*)). For comparison, a similar ensemble was constructed between lysin and VERL repeat 1 using a lysin solution structure (PDB 5utg) and a VERL repeat 1 crystal structure (PDB 5ii4) using constraints based on a cocrystal structure of lysin and VERL repeat 3 (PDB 5mr3). Figures of 3D protein models were produced using PyMOL (v.1.8, Schrodinger, LLC), regular secondary structure defined using the DSS function, and electrostatic surfaces calculated using the APBS/PDB2PQR server (*50, 51*).

*Characterization of FITZ complexes by dynamic light scattering*

Dynamic light scattering measurements were performed using a Zetasizer Nano (Malvern Pananalytical). Natural lysin purified from red abalone testis lysate was standardized to 100 μM in 10 mM Tris, pH 7.4, and light scattering was measured with and without equimolar FITZAP-8D at different salt concentrations (150 and 500 mM NaCl). Six technical replicates of 15 scans each were collected and averaged. FITZAP-8D in isolation produced no substantial light scattering over buffer.

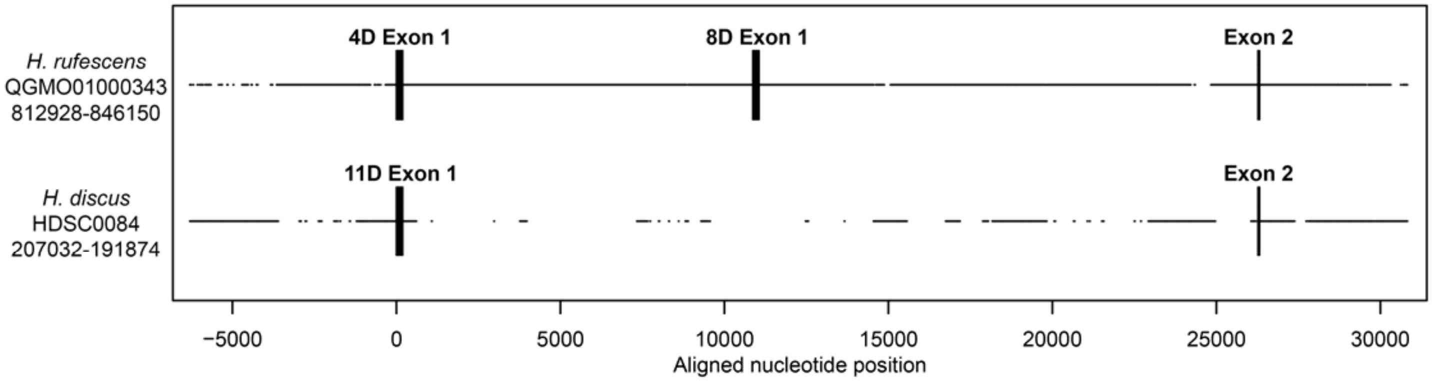


**Fig. S1. Genomic structure of FITZAP locus in red and disk abalone.** Comparison of FITZAP genomic loci between the red and disk abalone draft genomes. Each FITZAP coding sequencing is spliced using two exons: exon 1 contains the signal peptide, poly-aspartate array, lysin binding region, and the furin cleavage site, while exon 2 contains the majority of the FITZAP C-terminal peptide. The multiple FITZAP isoforms are produced through alternative splicing of optional exon 1 sequences, with the red FITZAP 4D exon 1 being orthologous to disk FITZAP-11D.

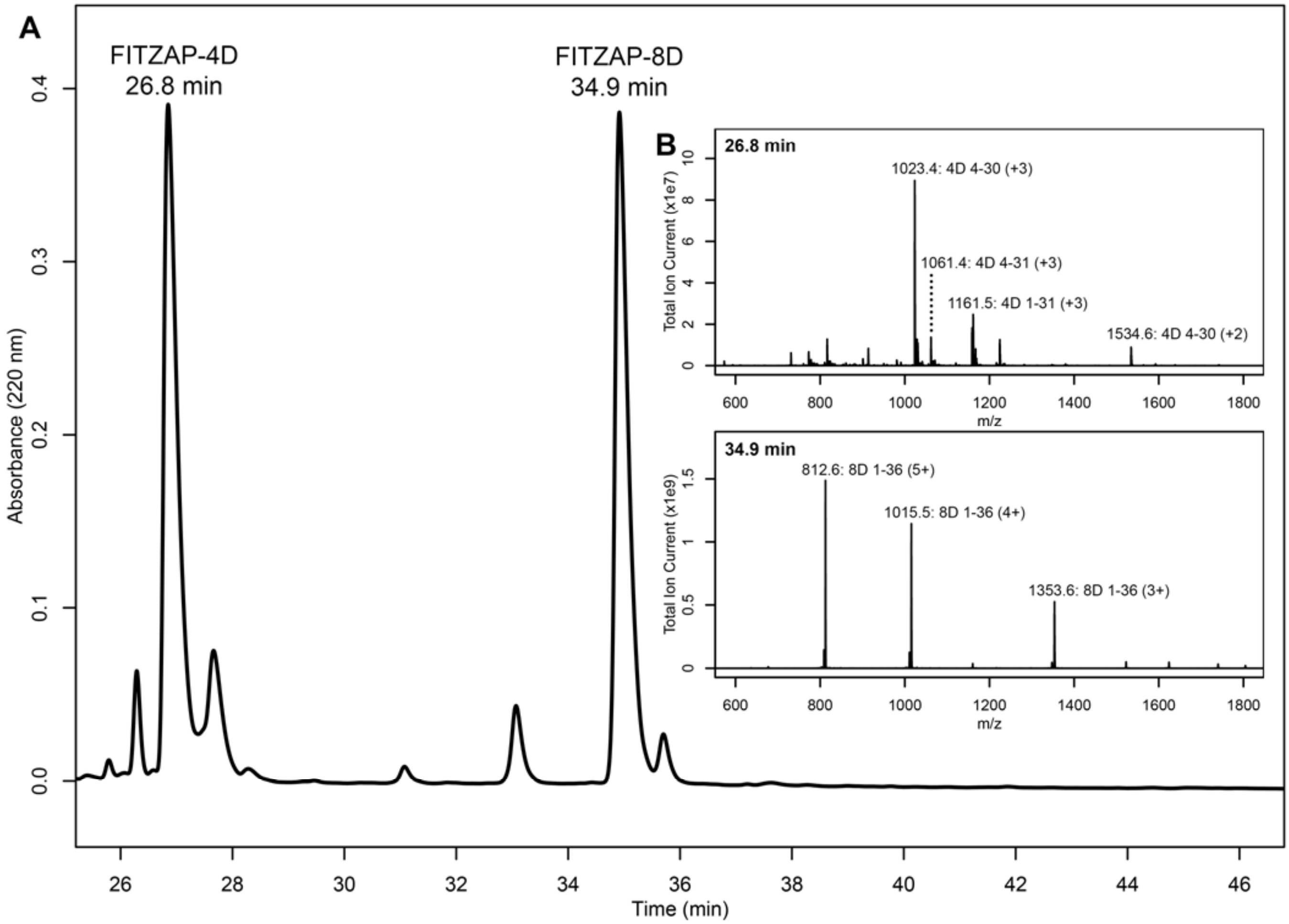


**Fig. S2. Purification and mass spectral analysis of natural red FITZAP-4D and 8D.** (A) Separation of anion exchange-enriched red abalone sperm lysate, with major peaks at 26.8 and 34.9 min being FITZAP-4D and 8D, respectively. (B) Mass spectral analysis of the FITZAP fractions show masses consistent with processing of FITZAP precursors by furin and carboxypeptidase B to liberate the C-terminal peptide from the longer N-terminal peptide containing the poly-aspartate region.

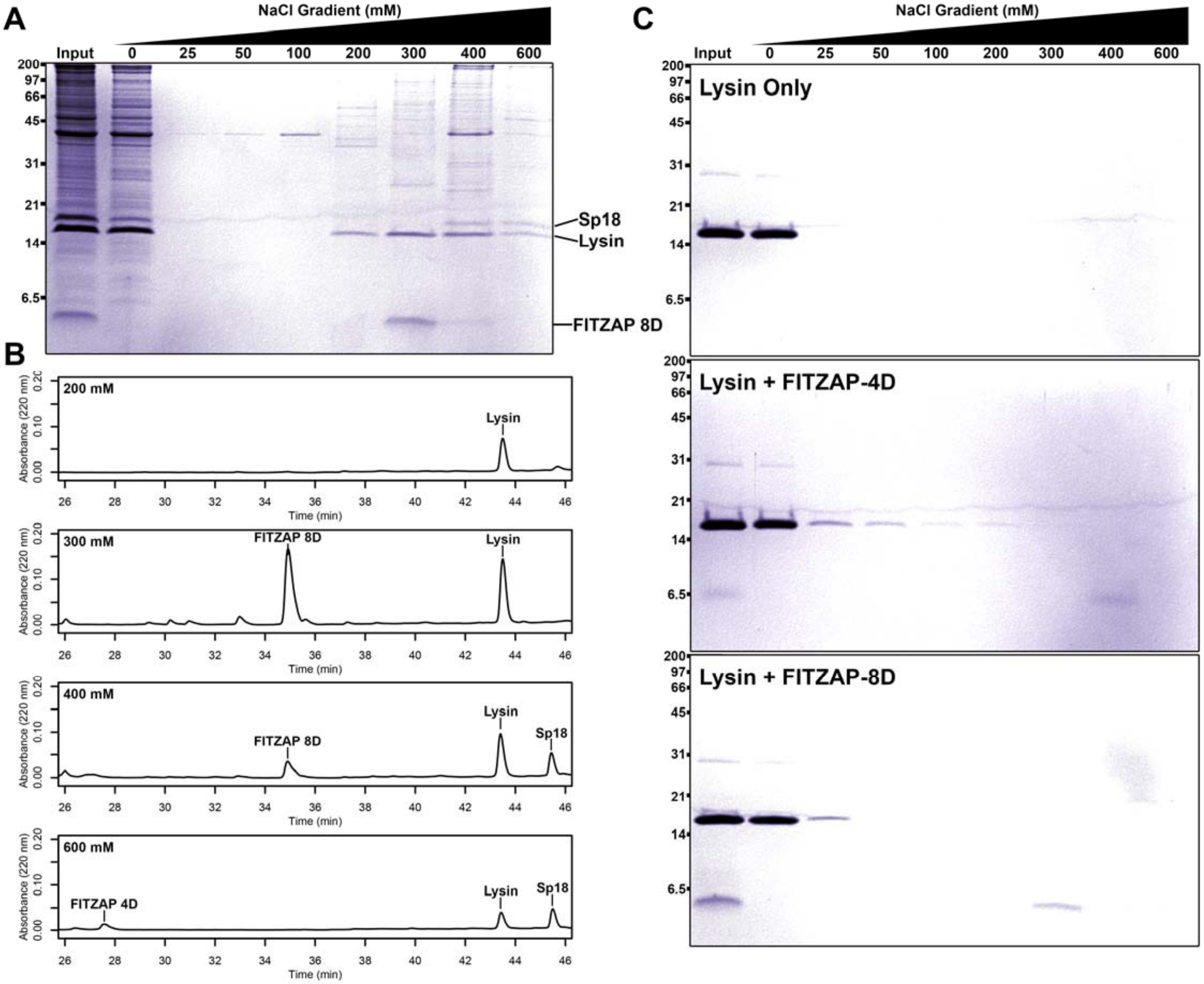


**Fig. S3. Co-purification of FITZAP and cationic acrosomal proteins by anion exchange.** (A) SDS-PAGE of proteins from red abalone sperm lysate separated by anion exchange. Despite their high net positive charges, lysin and sp18 were observed co-elute with FITZAP isoforms in the highest salt fractions. (B) RP-HPLC analysis of select anion exchange fractions which contained lysin. The peak of lysin elution at 300 mM coincides with the peak elution of FITZAP-8D, with both sp18 and FITZAP-4D eluting under higher salt conditions. (C) Purified lysin has no innate affinity for the anion exchange resin, yet its elution is retarded when mixed with either FITZAP isoform.

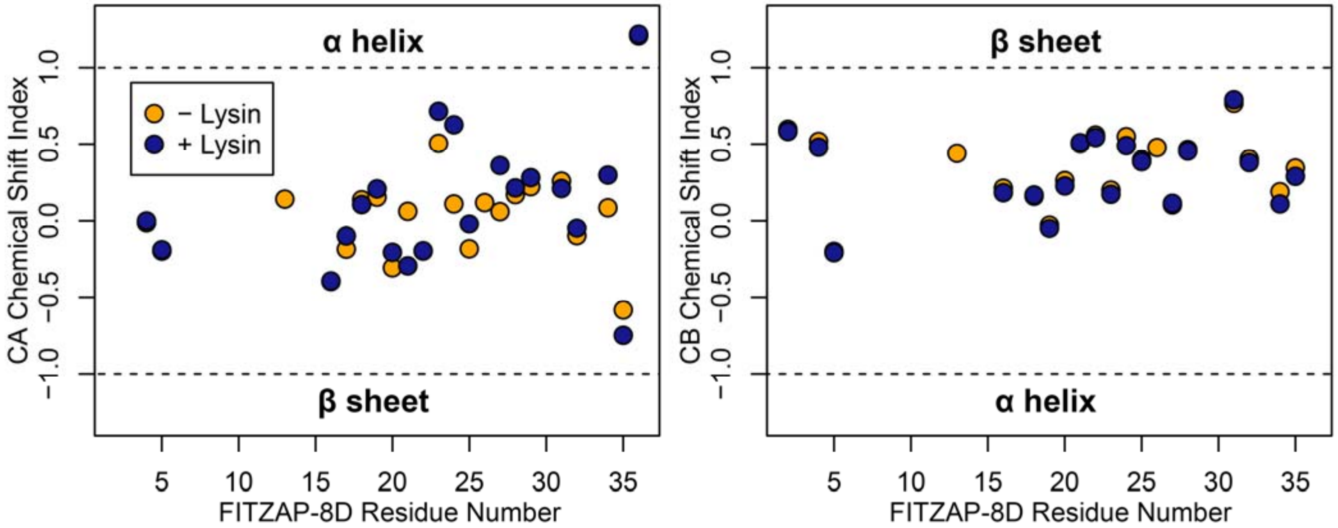


**Fig. S4. FITZAP is intrinsically disordered when free and bound to lysin.** CA and CB chemical shift indices for FITZAP-8D support that it is an intrinsically disordered protein, in both the presence and absence of equimolar lysin.

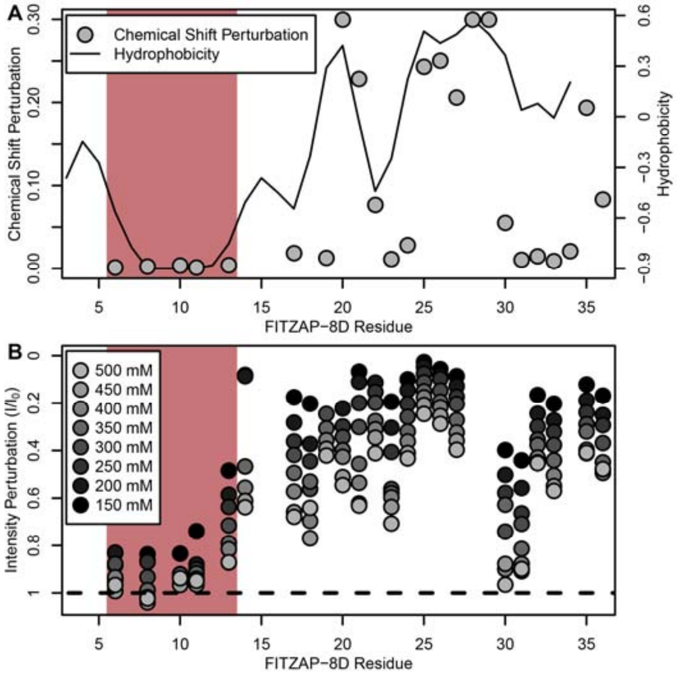


**Fig. S5. NMR perturbation of FITZAP-8D upon lysin binding.** (A) Backbone amide chemical shift perturbation of FITZAP-8D upon addition of lysin under seawater conditions (500 mM NaCl). The solid line denotes the relative hydrophobicity of the sequence which correlates well with the chemical shift perturbation, supporting that FITZAP-lysin interactions under high salt conditions are driven by hydrophobic packing. The poly-aspartate region is highlighted in red and shows no chemical shift perturbation under seawater conditions. (B) Intensity perturbation of FITZAP-8D by binding of lysin under different ionic strengths ranging from approximately intracellular levels (150 mM) to seawater conditions (500 mM). Differences in intensity perturbation as a function of salt are in part driven by the salt dependence on lysin-FITZAP $K_d$ (Fig 3), but the weak perturbation observed in the poly-aspartate region only under low salt conditions support that these residues are involved in some form of molecular interaction with lysin (likely salt bridges as part of FITZ complexes) only under intracellular conditions.

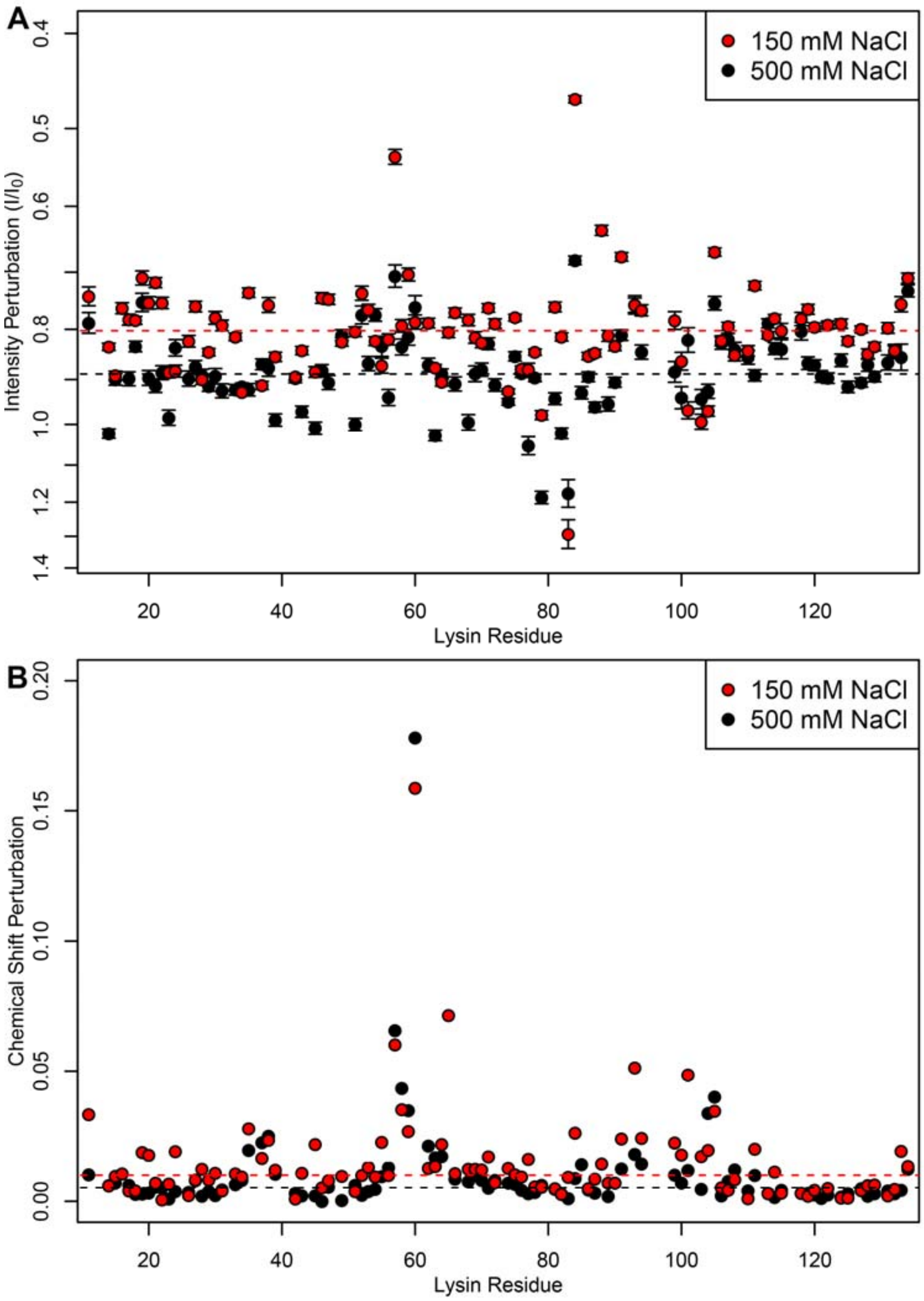


**Fig. S6. FITZ complexes experience different exchange rates in intracellular vs seawater conditions.** (A) Intensity perturbation and (B) chemical shift perturbation of lysin in the presence of FITZAP-8D (one molar equivalent) under both intracellular (150 mM) and seawater (500 mM) salt conditions, with dotted lines representing the median of each condition. Under intracellular salt concentrations, there is greater intensity perturbation, reflective of greater peak broadening and faster transverse relaxation from increased molecular weight of FITZAP binding lysin. However, despite lower affinity under higher salt concentrations, similar if not greater chemical shift perturbation is observed under seawater ionic strengths. Chemical shift perturbations are observed under fast exchange dynamics (ns-µs scales), while intensity perturbations imply slower dynamics (ms scales), such that this perturbation suggests that lysin-FITZAP are in slower exchange under intracellular salt conditions.

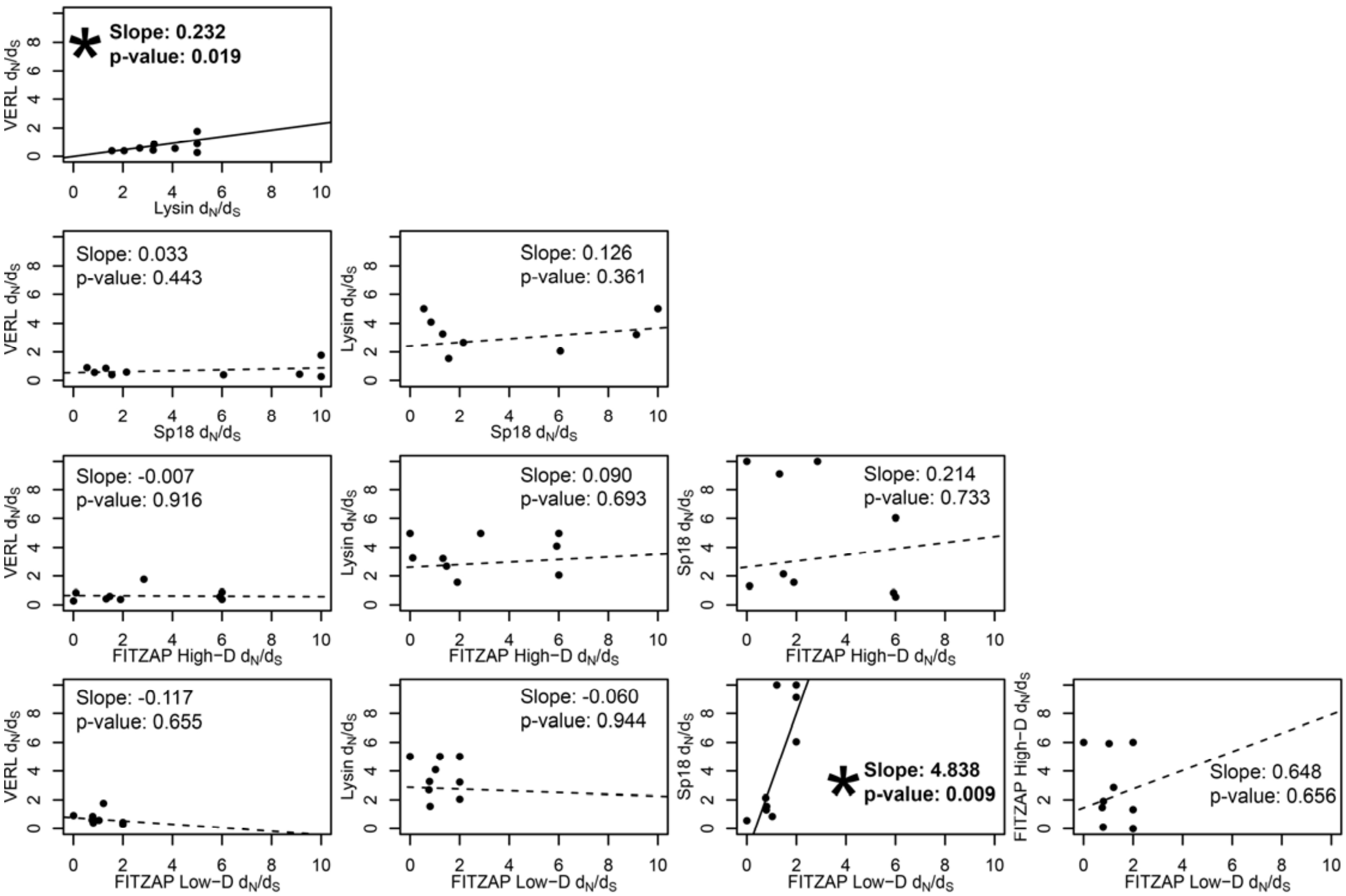


**Fig. S7. Test of sequence coevolution between abalone reproductive genes by correlated branch $d_N/d_S$.** Comparison of branch dN/dS estimates for all combinations of lysin, VERL, sp18, FITZAP low-D, and FITZAP high-D across the Pacific abalone clade. Based on methods by Clark et al. (*27*), correlation between these values was assessed by weighted linear regression, with a significant positive slope suggesting coevolution. Significant correlations were found between lysin-VERL (as in Clark et al. (*27*)) and in sp18-FITZAP low-D.
